# Late Pleistocene palaeoecology and phylogeography of woolly rhinoceroses

**DOI:** 10.1101/2021.01.08.425884

**Authors:** Alba Rey-Iglesia, Adrian M. Lister, Anthony J. Stuart, Hervé Bocherens, Paul Szpak, Eske Willerslev, Eline D. Lorenzen

## Abstract

The woolly rhinoceros (*Coelodonta antiquitatis*) was a cold-adapted herbivore, widely distributed from western Europe to north-east Siberia during the Late Pleistocene. Previous studies associate the extinction of the species ~14,000 years before present to climatic and vegetational changes, and suggest that later survival of populations in north-east Siberia may relate to the later persistence of open vegetation in that region. Here, we analyzed carbon (*δ*^13^C) and nitrogen (*δ*^15^N) stable isotopes and mitochondrial DNA sequences to elucidate the evolutionary ecology of the species. Our dataset comprised 286 woolly rhinoceros isotopic records, including 192 unpublished records, from across the species range, dating from >58,600 ^14^C years to ~14,000 years before present. Crucially, we present the first 71 isotopic records available to date of the 15,000 years preceding woolly rhinoceros extinction. The data reveal ecological flexibility and geographical variation in woolly rhinoceros stable isotope compositions through time. In north-east Siberia, we detected *δ*^15^N stability through time. This could reflect long-term environmental stability, and might have enabled the later survival of the species in the region. To further investigate the palaeoecology of woolly rhinoceroses, we compared their isotopic compositions with that of other contemporary herbivores. This analysis suggests possible niche partitioning between woolly rhinoceros and both horse (*Equus spp.*) and woolly mammoth (*Mammuthus primigenius*), and isotopic similarities between woolly rhinoceros and both musk ox (*Ovibos moschatus*) and saiga (*Saiga tatarica*) at different points in time. To provide phylogeographical context to the isotopic data, we analyzed 61 published mitochondrial control region sequences. The data show a lack of geographic structuring; we found three haplogroups with overlapping distributions, all of which show a signal of expansion during the Last Glacial Maximum. Furthermore, our genetic findings support the notion that environmental stability in Siberia had an impact on the paleoecology of woolly rhinoceroses in the region. Our study highlights the utility of combining stable isotopic records with ancient DNA to advance our knowledge of the evolutionary ecology of past populations and extinct species.

## 1. Introduction

The Late Pleistocene was characterized by large-scale climatic and environmental change (Hubberten 2004). Around 33 thousand years before present (ka BP), a progressive cooling started, which led to the cold and dry environment of the Last Glacial Maximum (LGM; ~28.6 - 20.5 ka BP). This was followed by an increase in temperature, which peaked at the start of the Holocene 11.7 ka BP. These pronounced climatic perturbations led to shifts in the geographic distribution of species and in the composition of entire ecosystems.

During the Late Pleistocene, large mammal species, including cave bear (*Ursus spelaeus),* cave lion (*Panthera spelaea*), and woolly rhinoceros (*Coelodonta antiquitatis*) went extinct in northern Eurasia (Pacher and Stuart 2009; Stuart and Lister 2011, 2012). Other Pleistocene megafauna, including giant deer (*Megaloceros giganteus*) and woolly mammoth (*Mammuthus primigenius*), experienced strong reductions in their distributions during the Late Pleistocene, but only disappeared from the faunal record later, during the Holocene (Stuart et al. 2002; Lister and Stuart 2019). These different extinction patterns among species have been attributed to the interplay of several drivers, including climatic, anthropogenic, and/or genetic factors (Cooper et al. 2015; Pečnerová et al. 2017).

In the northern hemisphere, Late Pleistocene glacial phases were characterized by the mammoth steppe ecosystem, which comprised a mosaic of steppe-tundra and shrub vegetation, with high nutrient soils that enabled the growth of plants that sustained grazing species (Guthrie 1982, 2001). The mammoth steppe extended from the northern Iberian Peninsula to Canada, and was characterized by a diverse community of large herbivores.

The woolly rhinoceros was one of the iconic inhabitants of this steppe-tundra ecosystem. The species was adapted to the cold environment, and was covered with thick and long hair, as documented by mummified remains, but is believed not to have been well adapted to snowfall, due to its massive body on short legs (Boeskorov et al. 2011). Teeth structure and mesowear analysis indicate a grazing diet (Stuart and Lister 2012; Rivals and Lister 2016; Stefaniak et al. 2020), which is supported by grass remains recovered between the teeth of some specimens (Guthrie 1990). Microwear analysis of North Sea woolly rhinoceros suggests periodical inclusion of woody components in their diet (Van Geel et al. 2019). Pollen analysis of stomach contents has primarily identified grasses and sagebrushes (e.g. Boeskorov et al. 2011), and genetic analysis of stomach and gut content supports a diet of primarily grasses, with a contribution of forbs (Willerslev et al. 2014).

Prior to their extinction ~14 thousand calendar years before present (cal kyr BP), woolly rhinoceroses ranged from western Europe to north-east Siberia (Figure 1). Specimens have not been recovered from certain areas in Europe and north-central Siberia, suggesting the species was absent from these regions (Stuart and Lister 2012). Despite their presence in the fossil record in far northeastern Siberia, adjacent to the Bering Land Bridge, woolly rhinoceroses did not colonise North America; it has been suggested that the environmental conditions of the Bering Land Bridge and the paleoecology of eastern Beringia were not adequate for the species (Boeskorov 2001; Stuart and Lister 2012).

**Figure 1.**
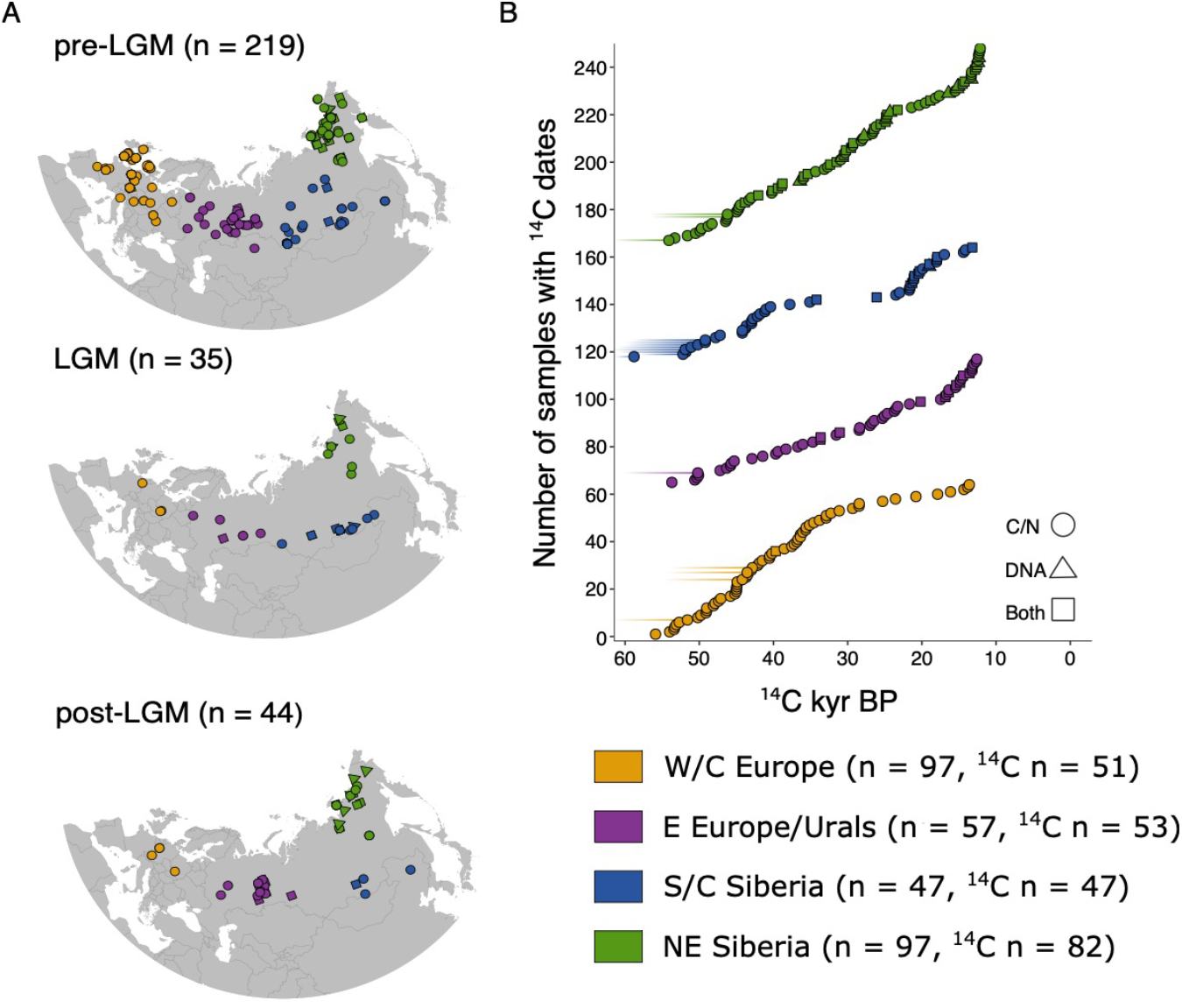
(A) Map of sample locations of 298 woolly rhinoceros remains analysed in this study, which were split into four geographic regions, indicated by different colours. Shapes represent data type: stable isotopes (circle), DNA (triangles), and both (square). Maps show three time bins: pre-LGM (>24,600 ^14^C years BP / 28,660 cal years BP), LGM (24,600 - 17,000 ^14^C years BP / 28,660 - 20,520 cal years BP), and post-LGM (<17,000 ^14^C years BP / < 20,520 cal years BP). (B) Chronology of the ^14^C radiocarbon dated samples used in this study for each region. Plotted dates represent uncal ^14^C years BP, extended lines represent infinite dates. Symbol colour and shape represents region and data type, respectively. Sample size (n = all samples; n = ^14^C dated samples) is indicated for each region.

Despite a progressive eastward contraction of the species’ range starting ~35 cal kyr BP (Stuart and Lister 2012), fossil evidence indicates woolly rhinoceroses were still present in large parts of Eurasia until ~16 cal kyr BP, suggesting an almost synchronous or at least rapid extinction across their range (Lorenzen et al. 2011). Demographic modelling based on genomic data has documented a population increase ~30 cal kyr BP, followed by demographic stability until close to woolly rhinoceros extinction (Lord et al. 2020). From 14.6 cal kyr BP, the range of the species had contracted such that woolly rhinoceroses were found only in the Ural Mountains, southern Siberia, and NE Siberia. The last dated fossil records are ~14 cal kyr BP from NE Siberia (Boeskorov 2001; Stuart and Lister 2012).

A combination of climate and vegetational change has been invoked as the main causative agent of woolly rhinoceros extinction (Lorenzen et al. 2011; Stuart and Lister 2012; Lord et al. 2020). These environmental shifts included increased ground moisture due to precipitation and snowfall in the Late Glacial Interstadial (14.7 - 12.9 cal kyr BP) (Sher 1997), and the spread of shrubs and trees (Stuart and Lister 2012). These new environmental conditions would have restricted areas of firm ground and open vegetation that are believed to have been the most suitable habitat for woolly rhinoceroses. The later survival of the woolly rhinoceros in NE Siberia may be linked to the late persistence of open vegetation in the region, compared with the rest of Eurasia (Stuart and Lister 2012). NE Siberia also played a key role for other Late Pleistocene megafauna species. When the Bering Land Bridge was exposed, it was used as a corridor to Eastern Beringia (e.g. by musk ox (*Ovibos moschaliis*) and woolly mammoth), and the region was the last mainland holdout of both steppe bison (*Bison priscus*) and woolly mammoth prior to their extinctions (Vartanyan et al. 1993; Kirillova et al. 2015a).

Stable isotope (*δ*^13^C and *δ*^15^N) analyses of bone and tooth collagen have provided important insights into the diet and palaeoecology of Late Pleistocene megafauna, including brown bear (*Ursus arctos*), cave bear, cave lion, giant rhinoceros (*Elasmotherium sibiricum*), musk ox, saiga (*Saiga tatarica*), and woolly mammoth (Szpak et al. 2010; Bocherens et al. 2011; Raghavan et al. 2014; Krajcarz et al. 2016; Jürgensen et al. 2017; Arppe et al. 2019; Kosintsev et al. 2019; Rey-Iglesia et al. 2019). *δ^13^C* and *δ*^15^N have also been used to investigate the paleoecology and paleodiet of the woolly rhinoceros (Bocherens et al. 1995, 1996, 1997, 2005, 2011; Higham et al. 2006; Jacobi et al. 2006, 2007, 2009; Kirillova et al. 2015b; Yates et al. 2017; Stefaniak et al. 2020). However, in contrast with work on e.g. woolly mammoth and saiga (Szpak et al. 2010, Jürgensen et al. 2017), a comprehensive, range-wide study of woolly rhinoceros paleoecology and paleodiet based on stable isotope data has so far been lacking.

In herbivores, *δ*^13^C and *δ*^15^N values reflect the isotopic compositions of plants in their diet (DeNiro 1985; Bocherens and Drucker 2007), the composition of which in turn is influenced by climatic and environmental factors (Hartman 2011; Murphy et al. 2013; Bonafini et al. 2013). Carbon isotopic compositions in herbivore bone and tooth collagen reflect the photosynthetic pathways and environmental parameters of plants at the base of the food web (Bocherens 2003). Nitrogen derives from the plant protein the animal digests; dietary choices may lead to differences in *δ*^15^N, even at the same trophic level (Bocherens 2003). Furthermore, environmental factors such as water stress, salinity or grazing pressure, are known to affect plant *δ*^15^N values, and changes in vegetation and soil nutrient cycling will therefore be recorded in bone and tooth collagen.

In this study, we used *δ*^13^C and *δ*^15^N retrieved from woolly rhinoceros remains spanning 45,000 years, to investigate Late Pleistocene changes in palaeoecology and population history. We present 192 new isotopic records from across the species range dating from >58,600 ^14^C years to 14,140 cal yr BP. These include 71 isotopic records from the 15 kyr preceding woolly rhinoceros extinction, which represent the first available stable isotope records from this period. We combined the 192 new records with published *δ*^13^C and *δ*^15^N data of woolly rhinoceros, totalling 286 samples, and with comparative data from five other contemporaneous megaherbivore species (horse (*Equus spp.*), musk ox, reindeer (*Rangifer tarandus*), saiga, woolly mammoth) to (i) investigate temporal and spatial changes in woolly rhinoceros diet and ecology; (ii) specifically assess changes in the millennia preceding extinction; (iii) elucidate the late survival of the species in NE Siberia; (iv) understand the broader ecological context of our findings. In addition, to investigate the phylogeographical context of the palaeoecological data, we included the analysis of a comprehensive data set of available mitochondrial DNA (mtDNA) control region sequences.

## 2. Material and methods

### 2.1 Stable isotope data

We present 192 new *δ*^13^C and *δ*^15^N measurements from woolly rhinoceros (Supplementary Table 1), determined by isotope ratio mass spectrometry (IRMS); ^14^C radiocarbon dates for these samples were published in previous studies (Lorenzen et al. 2011; Stuart and Lister 2012). The samples ranged in age from infinite to 14,140 cal yr BP. The spatial distribution of the material spans northern Eurasia, with records from Spain to NE Siberia (Figure 1). Radiocarbon dating and isotopic measurements of samples were conducted at Oxford Radiocarbon Accelerator Unit (University of Oxford) and Aarhus AMS Centre (University of Aarhus). Reliability of the stable isotopic values was established by measuring the atomic C:N ratio. All our samples fell within the accepted range of 2.9 to 3.6 (DeNiro 1985; Ambrose 1990).

We furthermore compiled a comprehensive database of available *δ*^13^C and *δ*^15^N stable isotope records for woolly rhinoceros (Supplementary Table 1). We used Google Scholar to perform a literature search with the terms “woolly rhinoceros” and “stable isotopes”, and *“Coelodonta antiquitatis”* and “stable isotopes”. We recovered data for 94 specimens, derived from ten published studies and spanning the species’ range (Bocherens et al. 1995, 1996, 1997, 2005, 2011; Higham et al. 2006; Jacobi et al. 2006, 2007, 2009; Kirillova et al. 2015b; Kuc et al. 2012; Yates et al. 2017). All previously published data represented samples > 28,660 cal yr BP. Our final dataset represented 286 samples (Figure 1).

To contextualize the woolly rhinoceros isotopic data with other coeval large herbivores, we performed a literature search and compiled *δ*^13^C and *δ*^15^N for five other mammoth steppe species from Eurasia: horse, musk ox, reindeer, saiga, and woolly mammoth (Supplementary Table 2). Paleoecological inference based on *δ*^13^C and *δ*^15^N data is highly influenced by regional environmental characteristics, and large isotopic differences have been observed for *δ*^13^C and *δ*^15^N between the North American and Eurasian portions of the mammoth steppe (Szpak et al. 2010). As woolly rhinoceroses were only present in Eurasia, we omitted North American records for the other species.

### 2.2 Stable isotope analysis

We recalibrated the original ^14^C dates using OxCal 4.3 (Ramsey 2009) and the IntCal20 (Reimer et al. 2020) calibration curve. Unless mentioned, dates are reported as calibrated years before present (cal yr BP). Dates reported as infinite correspond to samples where the ^14^C content was too small to produce a finite radiocarbon age.

To investigate temporal variation in the isotopic signature of woolly rhinoceroses and other megaherbivores, we divided the isotopic records into three time bins (Figure 1), following Kuitems et al. 2019: pre-LGM (> 24,600 ^14^C yr BP / > 28,660 cal yr BP), LGM (24,600 - 17,000 ^14^C yr BP / 28,660 - 20,520 cal yr BP), and post-LGM (< 17,000 ^14^C yr BP / < 20,520 cal yr BP). The post-LGM cut-off date corresponds to 12,140 ^14^C yr BP or 14,140 cal yr BP, the age of the youngest woolly rhinoceros specimen in our dataset. The sample size of each time bin was: pre-LGM n = 215, LGM n = 32, post-LGM n = 39 (Table 1).

**Table 1.** Summary of *δ*^13^C and *δ*^15^N values of the 286 woolly rhinoceros specimens used in this study. Number of specimens in each temporal/spatial bin is indicated in parenthesis.

| Time bin | Region (sample size) | $\delta^{13}\text{C}$ | | | | | $\delta^{15}\text{N}$ | | | | |
| --- | --- | --- | --- | --- | --- | --- | --- | --- | --- | --- | --- |
|  |  | Min | Max | Median | Mean | sd | Min | Max | Median | Mean | sd |
| pre-LGM (>24,600 BP / 28,660 cal BP) | W/C Europe (90) | -22.5 | -18.0 | -20.0 | -20.0 | 0.7 | 0.7 | 11.5 | 5.3 | 5.3 | 1.8 |
|  | E Europe/Urals (34) | -20.8 | -18.6 | -19.5 | -19.5 | 0.5 | 2.6 | 9.8 | 5.1 | 5.4 | 1.8 |
|  | S/C Siberia (26) | -20.5 | -18.5 | -19.6 | -19.6 | 0.6 | 3.0 | 9.1 | 6.6 | 6.1 | 1.5 |
|  | NE Siberia (65) | -22.7 | -19.5 | -20.6 | -20.6 | 0.6 | 2.6 | 10.8 | 6.6 | 6.7 | 1.9 |
| LGM (24,600-17,000 BP / 28,660-20,520 cal BP) | W/C Europe (3) | -20.0 | -19.0 | -20.0 | -19.7 | 0.6 | 3.0 | 5.0 | 4.0 | 4.0 | 1.0 |
|  | E Europe/Urals (6) | -19.6 | -19.0 | -19.0 | -19.1 | 0.2 | 4.0 | 8.8 | 5.0 | 5.4 | 1.8 |
|  | S/C Siberia (16) | -20.1 | -18.4 | -19.1 | -19.1 | 0.5 | 3.1 | 10.3 | 4.9 | 5.2 | 1.8 |
|  | NE Siberia (7) | -20.7 | -19.3 | -20.1 | -20.0 | 0.6 | 2.8 | 10.6 | 6.8 | 6.2 | 2.6 |
| post-LGM (<17,000 BP / < 20,520 cal BP to extinction) | W/C Europe (4) | -20.1 | -20.0 | -20.0 | -20.0 | 0.1 | 3.0 | 4.0 | 3.1 | 3.3 | 0.5 |
|  | E Europe/Urals (17) | -20.6 | -19.0 | -20.0 | -19.8 | 0.5 | 1.9 | 11.4 | 3.2 | 3.9 | 2.4 |
|  | S/C Siberia (4) | -20.4 | -18.6 | -19.8 | -19.7 | 0.9 | 4.2 | 8.2 | 4.8 | 5.5 | 1.9 |
|  | NE Siberia (14) | -22.4 | -19.7 | -20.2 | -20.3 | 0.7 | 2.3 | 9.5 | 6.7 | 6.3 | 2.7 |

To accommodate for differences in fossil chronology across Eurasia (such as the later survival of woolly rhinoceros in NE Siberia) and the environmental heterogeneity of Eurasia during the Late Pleistocene, we divided the data into four geographical regions: (i) western and central Europe (n = 97); (ii) European Russia, Urals and Kazakhstan (n = 57); (iii) southern and central Siberia and China (n = 46); and (iv) north-east Siberia (n = 86, Table 1). To simplify, we refer to these regions as W/C Europe, E Europe/Urals, S/C Siberia, and NE Siberia, respectively.

Stable isotopes were retrieved from both bone and tooth. The isotopic signature in bone material provides information on the average diet of an individual in the years preceding death (Hedges et al. 2007; Hobson and Clark 1992). In contrast, data from mammalian tooth collagen corresponds to the early years of an individual’s life, depending on the chronology of tooth development for the species in question, and their *δ*^15^N signature may reflect the suckling ^15^N-enriched composition (DeNiro 1985; Fizet et al. 1995; Bocherens and Drucker 2007). For the purposes of our analysis, we assumed that the stable isotope composition of bone and tooth collagen are comparable, albeit systematic differences in *δ*^13^C and *δ*^15^N isotopic abundances between bone and dentine have been recognized in some mammals. For instance, lower *δ*^15^N values in bone compared to dentine have been measured in carnivores such as bears and hyenas (Bocherens et al. 1997; Bocherens 2015), or in herbivores such as reindeer (Fizet et al. 1995). For the species where the isotopic differences between bone and tooth have been directly identified and quantified, *δ*^13^C and *δ*^15^N data are adjusted before comparing bone and tooth isotopic values. However, there has been no systematic study characterising possible *δ*^13^C and *δ*^15^N isotopic differences between bone and tooth in woolly rhinoceros, and our data have thus not been corrected. We are aware of the bias that this might introduce to our data. However, assuming the correction factor for woolly rhinoceros is similar to estimates from other herbivore species, such as reindeer (*δ*^13^C = 0.2‰ - 0.5‰ and *δ*^15^N = 0.7‰ - 2‰, lower in bone than in dentine, Fizet et al. 1995), the skeletal element variation in our data set is less than the variation we detect among individuals across time bins and regions.

Groups (representing time or space) with low sample size (n < 30) were tested for normality using the Shapiro-Wilkinson test. As the data distribution for some of these groups was not normal, we chose to use non-parametric testing for all analyses comparing isotopic signatures for different time bins and regions. The Kruskal-Wallis test was used to determine statistical differences among regions for each time bin.

To further detect differences between pairs of time bins and/or regions, we used the post-hoc Kruskal Conover test applying the Bonferroni correction to control for Type I errors. Statistical data analysis was conducted in RStudio (Team RStudio 2018) using the packages nortest (Gross and Ligges 2015) and PMCMR (Pohlert 2018). Significant differences were reported at the 95% confidence level or p-value of 0.05.

Bivariate plots and boxplots were performed in RStudio (Team RStudio 2018) using the package ggplot2 (Wickham 2011). Summary statistics were obtained in RStudio.

Multispecies comparisons were also performed using SIBER (Jackson et al. 2011), this R package was used to estimate and compare the Standard Ellipse Area (SEA) as a proxy for isotopic niche range.

### 2.3 DNA data compilation and analysis

To contextualize the stable isotope data and evaluate genetic changes across time and space, we compiled all available ancient mtDNA control region sequences of woolly rhinoceros published to date (Lorenzen et al. 2011; Lord et al. 2020). Due to a large amount of missing data for the control region locus, eight of the 14 sequences from Lord et al. 2020 were not included in our analysis (Supplementary Table 3). We did not include the mtDNA sequence published in Willerslev et al. (2009), as the sample was not dated. The final dataset comprised 61 mtDNA control region sequences; 35 of these samples were also represented in the stable isotope analysis (Supplementary Table 3). Sequences were aligned in Geneious (Kearse et al. 2012), and the length of the final alignment used in our analysis was 541 basepairs (bp). We generated a haplotype network using the median-joining (Bandelt et al. 1999) algorithm implemented in the program PopART v1.7 (Leigh and Bryant 2015).

To estimate a phylogeny, we used BEAST v.1.10.4 (Drummond et al. 2012) using the radiocarbon dates of the sequences to estimate a mutation rate and divergence times. This analysis was performed using the same dataset as for the haplotype network. We used a coalescent constant size model, as data were from a single species, and applied a strict clock. The clock rate was set to a normal distribution with an initial value of 6.1×10^-9^ substitutions/site/year, a mean value of 6.1×10^-9^, and a standard deviation of 0.01 as in Lord et al. 2020. As estimated in PartitionFinder (Lanfear et al. 2012), we used the model HKY + I + G for all the codons of the control region.

All BEAST analyses were run in two independent MCMC chains of 1×10^7^generations each, sampling trees and model parameters every 1×10^3^ generations. Tracer v1.6 (Rambaut et al. 2018) was used to combine and inspect the results of each run and to determine the convergence of each parameter, all of which had ESS values > 200. We identified the Maximum Clade Credibility (MCC) tree in TreeAnnotator v1.8.0, and visualized and graphically edited the MCC tree using Figtree.

The covariation of stable isotopes and haplogroups was also explored. A bivariate plot was performed in RStudio (Team RStudio 2018) using the package ggplot2 (Wickham 2011). A cluster dendrogram was generated using the R package cluster (Maechler 2019). For this, we applied the Euclidean distance method algorithm with Ward’s minimum variance criterion to minimize the total within-cluster variance. We set the number of clusters to three, equivalent to the number of haplogroups recovered in the phylogenetic analysis.

## 3. Results

### 3.1 Stable isotope analysis: woolly rhinoceros

Summary statistics of woolly rhinoceros stable isotope measurements were compiled in Table 1 and visually explored in Figures 2 - 4. *δ*^13^C values ranged from −22.7‰ to −18.1‰, with an average value of −19.6 ± 0.71‰. *δ*^15^N values ranged from +0.7‰ to +11.5‰, with an average value of +5.7 ± 2.04‰. Summary statistics indicated a smaller range of variation for *δ*^13^C than *δ*^15^N.

**Figure 2.**
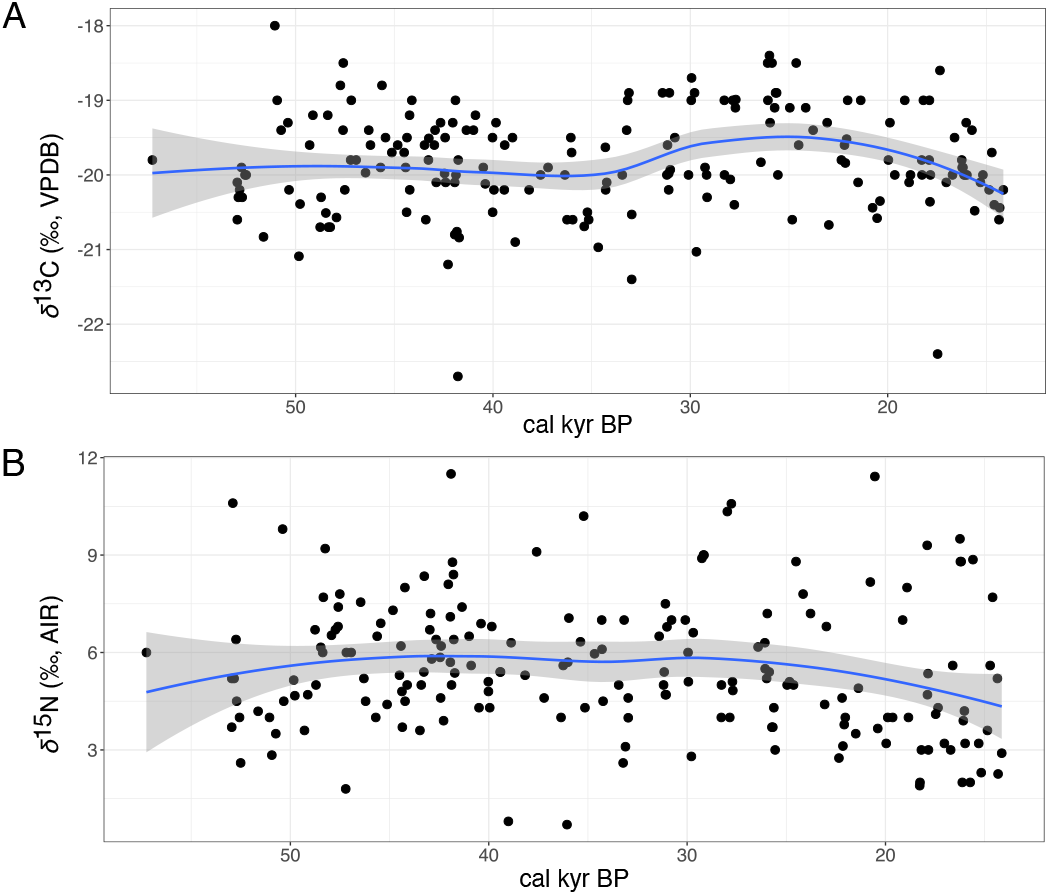
Temporal variation in (A) *δ*^13^C and (B) *δ*^15^N isotopic values across the 286 woolly rhinoceros sampled for this analysis. Local regression curves were estimated using the LOESS (locally estimated scatterplot smoothing) method as implemented in ggplot2 (Wickham 2011).

Figure 2A shows variation in *δ*^13^C through time. We detected an increase in *δ*^13^C starting ~34 cal kyr BP, which peaked at the onset of the LGM ~27 cal kyr BP, with a subsequent decline. We further explored differences in *δ*^13^C within the three time bins using bivariate plots, and found higher average *δ*^13^C during the LGM than for the other two time bins (Figure 3). Statistical testing showed significant differences in average *δ*^13^C between pre-LGM and LGM (*p* < 0.001), and between LGM and post-LGM (*p* = 0.003).

**Figure 3.**
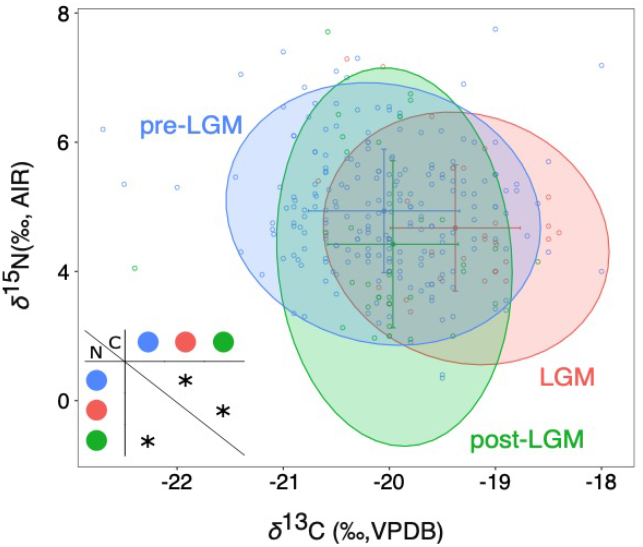
Bivariate plot of *δ*^13^C and *δ*^15^N isotopic values across 286 woolly rhinoceros specimens. Data were divided into three time bins: pre-LGM (n = 215), LGM (n = 32), and post-LGM (n = 39). Ellipses represent 0.95 confidence levels. Crosses represent mean value +/− s.d. for each time bin. Matrix represents pairwise comparison of differences in *δ*^13^C (below diagonal) and *δ*^15^N (above diagonal). Significant differences between time bins are indicated with an asterisk.

We investigated the influence of time *δ*^13^C within each of the four geographic regions (Figure 4A). For three regions (excluding W/C Europe), we detected significant changes in average *δ*^13^C over time (p-values: E Europe/Urals: *p* = 0.02; S/C Siberia: *p* = 0.03; NE Siberia: *p* = 0.008), with higher values during the LGM. Within each time bin, we identified differences in isotopic values across regions (Figure 4A, Supplementary Table 4).

**Figure 4.**
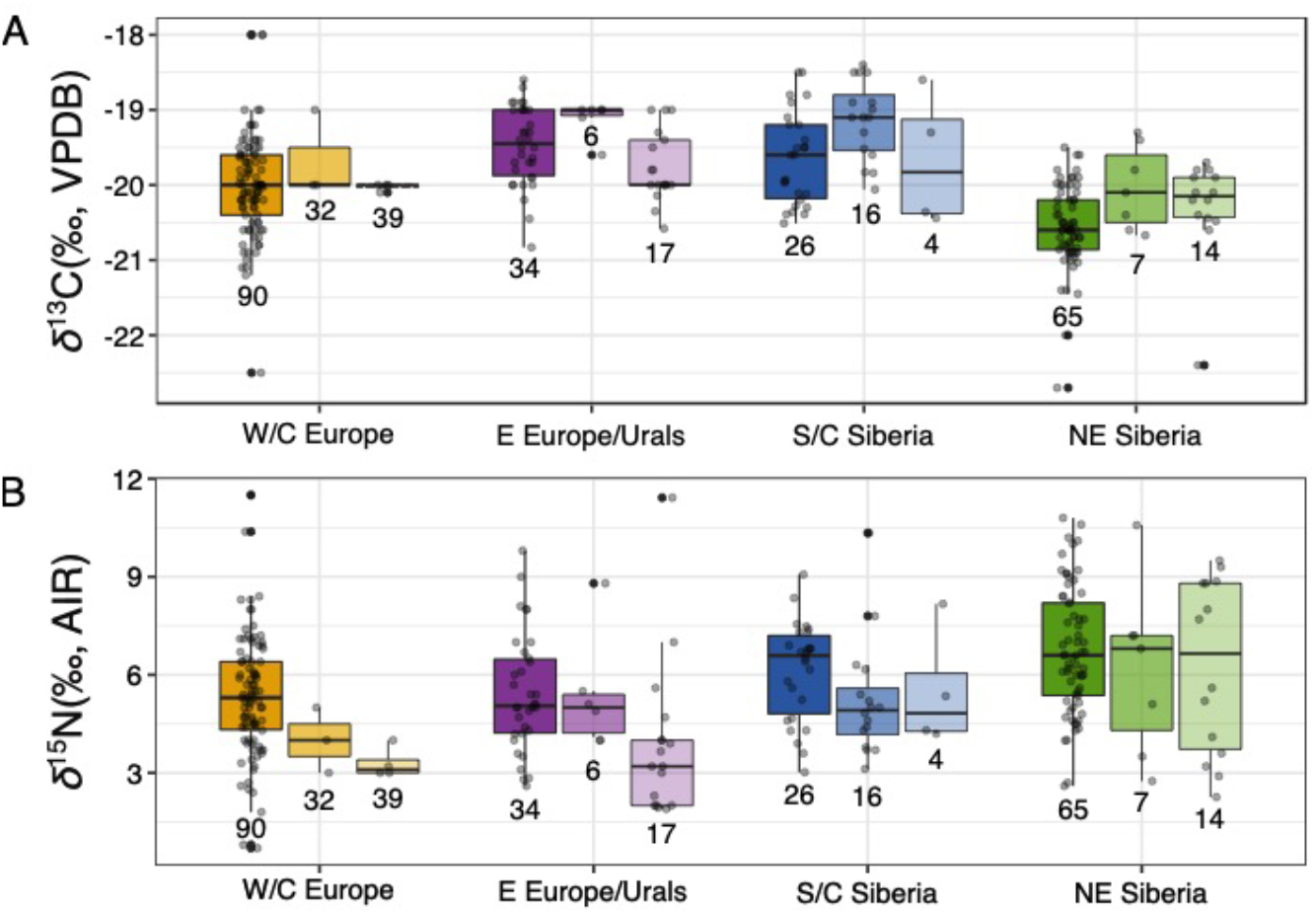
Boxplots showing spatial and temporal variation in (A) *δ*^13^C and (B) *δ*^15^N in woolly rhinoceroses. For each region, boxplots are organized in time bins from left to right: pre-LGM, LGM, and post-LGM. Sample sizes are indicated below each box plot. Box plots represent median, upper, and lower quartile, and whiskers represent highest and lowest value

For *δ*^15^N, we observed a gradual decline starting at the onset of the LGM ~27 cal kyr BP (Figure 2B). When we split the range-wide dataset into three time bins, statistical testing showed significant differences in *δ*^15^N between pre-LGM and post-LGM (*p* = 0.002) (Figure 3). When the data were further investigated across time for each region, we detected significant continuous declines in average *δ*^15^N over time for two regions (p-values: W/C Europe = 0.010, E Europe/Urals = 0.004), but not for S/C and NE Siberia (Figure 4B).

For pre-LGM, pairwise comparisons between regions showed significant differences between NE Siberia and both W/C Europe and E Europe/Urals (Figure 4B, Supplementary Table 4). For LGM, we did not identify any differences in *δ*^15^N between regions. For post-LGM, NE Siberia *δ*^15^N differed significantly from E Europe/Urals.

### 3.2 Stable isotope analysis: other herbivores

To contextualise woolly rhinoceros isotopic variation, we compared the data with stable isotope records from five co-distributed herbivore species. To make the data directly comparable across time and space, we restricted the geographical distribution of the data to Eurasia (Supplementary Table 2). Bivariate and boxplots of the data divided by species and by time bin correspond to Figures 5 and 6. Graphical exploration and statistical testing indicated that pre-LGM *δ*^13^C and *δ*^15^N of rhinoceros differed from all the other species surveyed (Figures 5-6, Supplementary Table 5). Rhinoceros LGM *δ*^13^C and *δ*^15^N were similar to saiga. Post-LGM rhinoceros *δ*^13^C and *δ*^15^N values were similar to musk ox.

**Figure 5.**
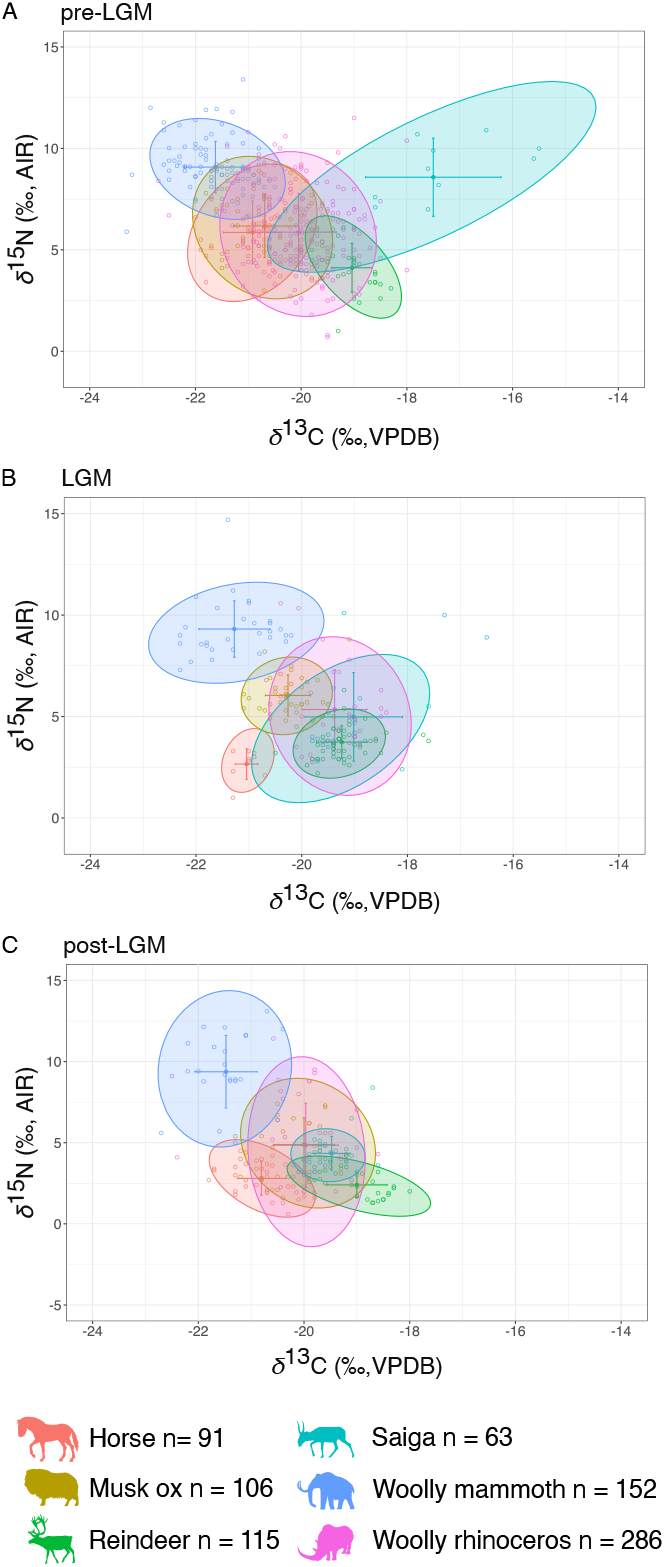
Bivariate plot of *δ*^13^C and *δ*^15^N isotopic composition of woolly rhinoceroses and other contemporary megaherbivores from Eurasia. Ellipses represent 0.95 confidence levels. Crosses represent mean value +/− s.d. for each species. Panels represent three time bins: (A) pre-LGM (>24,600 ^14^C years BP / 28,660 cal years BP), (B) LGM (24,600 - 17,000 ^14^C years BP / 28,660 - 20,520 cal years BP), and (C) post-LGM (<17,000 ^14^C yearsBP / < 20,520 cal years BP). Sample sizes for each species are indicated next to each silhouette. Horse, musk ox, reindeer, and woolly rhinoceros silhouettes: license Public Domain Dedication 1.0. Saiga silhouette: Andrey Giljov, license CC-BY-SA-4.0 (https://creativecommons.org/licenses/by-sa/4.0/deed.en). Woolly mammoth silhouette: Zimices, license CC BY-NC-3.0 (https://creativecommons.org/licenses/by-nc/3.0/).

**Figure 6.**
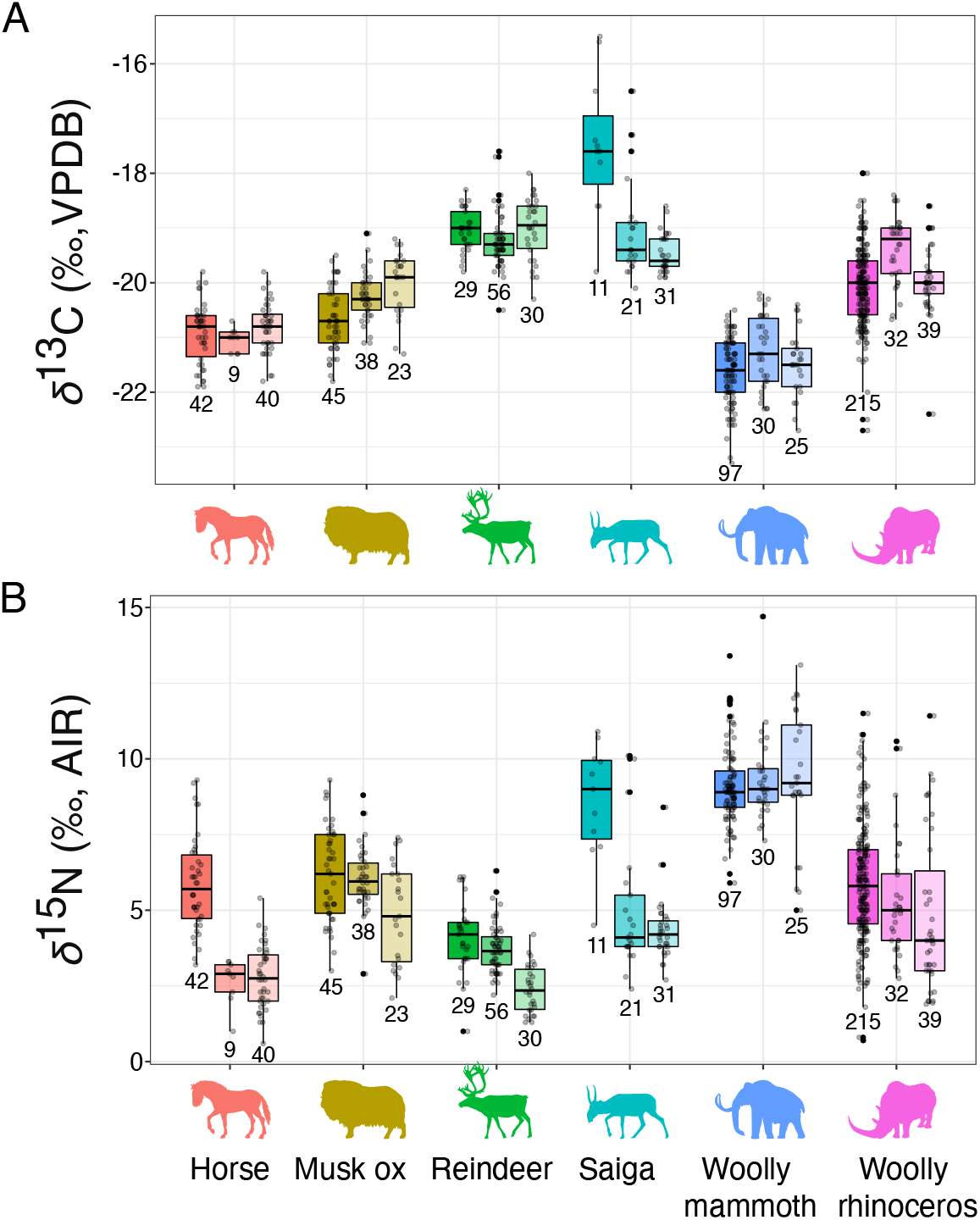
Boxplots representing isotopic variation over time for (A) *δ*^13^C and (B) *δ*^15^N in six Late Pleistocene megaherbivores. For each species, boxplots are organized from left to right: pre-LGM, LGM, and post-LGM. Sample sizes are indicated below each box plot. Box plots represent median, upper, and lower quartile, and whiskers represent highest and lowest value. Horse, musk ox, reindeer, and woolly rhinoceros silhouettes: license Public Domain Dedication 1.0. Saiga silhouette: Andrey Giljov, license CC-BY-SA-4.0 (https://creativecommons.org/licenses/by-sa/4.0/deed.en). Woolly mammoth silhouette: Zimices, license CC BY-NC-3.0 (https://creativecommons.org/licenses/by-nc/3.0/)

To estimate species isotopic ranges, we used the Standard Ellipse Area (SEA) function implemented in SIBER (Supplementary Table 6). For pre-LGM, saiga had the largest isotopic range (SEA = 5.16), followed by rhinoceros (SEA = 4.25). For LGM, we observed a decline in isotopic range for four species, except mammoth (SEA = 2.91) and saiga (SEA = 5.30). For rhinoceros, SEA was reduced to 3.65, still remaining second largest after saiga. For post-LGM, rhinoceros had the largest SEA (SEA = 4.98), with an increase in the ellipse area from the LGM (LGM SEA = 3.65). For rhinoceros, post-LGM isotopic range recovered to pre-LGM values, albeit with overall lower *δ*^15^N values.

### 3.3 DNA analysis

We identified three haplogroups (A-C) in the mtDNA control region data (Figure 7, Supplementary Figure 1, and Supplementary Table 3). The alignment contained 66 segregating sites; haplogroups A and B differed from each other in two nucleotide positions, and haplogroups B to C in six nucleotide positions.

**Figure 7.**
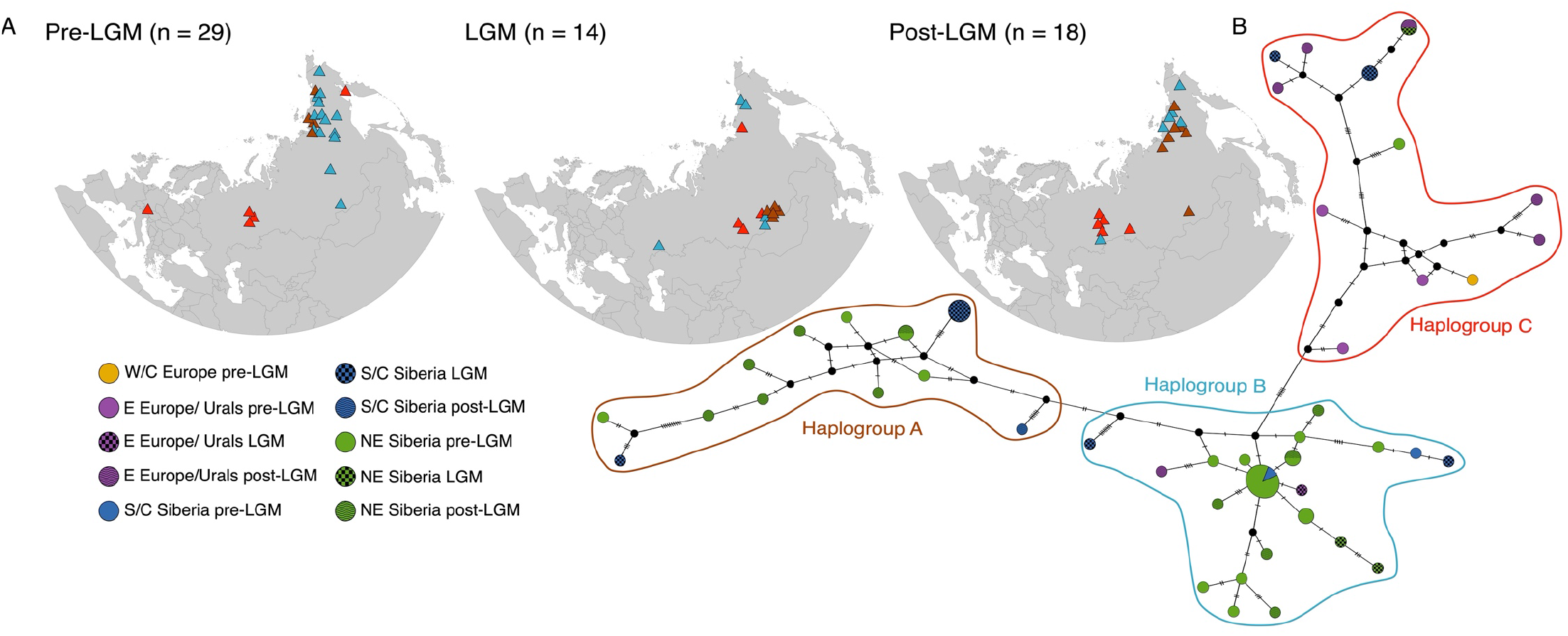
Mitochondrial haplotype network of 61 published control region sequences. (A) Geographical distribution of haplotypes across three time bins: pre-LGM, LGM, and post-LGM. Colours correspond to haplogroups, shown as polygons in panel B. (B) Haplotype network. Circles correspond to haplotypes, colours correspond to geographic origin of samples, and pattern corresponds to time bin (see legend). Circle size indicates number of samples per haplotype.

Haplogroup A was only found in S/C and NE Siberia; it was present in NE Siberia pre-LGM, in S/C Siberia during LGM, and in both regions post-LGM.

Haplogroup B extended from southern European Russia to NE Siberia: pre-LGM, haplogroup B was present in S/C and NE Siberia; during the LGM, we identified haplogroup B in samples from southern European Russia, S/C Siberia, and NE Siberia; post-LGM, the haplogroup was present in E Europe/Urals and NE Siberia.

Haplogroup C was present across the species range in pre-LGM, extending from central Europe to NE Siberia. During LGM, haplogroup C remained in C/S and NE Siberia. Post-LGM, only woolly rhinoceroses from E Europe/Urals carried this haplogroup.

We also explored the covariation of stable isotopes and haplogroups. However, we could not identify any relationship between them (Supplementary Figure 2).

## 4. Discussion

We analysed *δ*^13^C and *δ*^15^N stable isotope data from 286 woolly rhinoceros specimens sampled across Eurasia, spanning over 45,000 years of evolutionary history. Our analyses revealed variation across geographic regions; S/C and NE Siberia showed stability in isotopic composition across time, in contrast with other regions, which experienced variation in *δ*^15^N. Our analysis of 61 mtDNA sequences did not show any geographic structuring, but we recovered three well-differentiated haplogroups with overlapping distributions, all of which showed a signal of expansion during the LGM.

### 4.1 Temporal variation in *δ*^13^C within regions

Across samples, we found an average *δ*^13^C value of −19.9 ¼, indicating that woolly rhinoceroses had a diet dominated by C3 plants (Bocherens 2003); this is expected in herbivores from cold and temperate climates. We detected significant changes in average *δ*^13^C across time bins; between pre-LGM and LGM, and between LGM and post-LGM (Figure 3).

Although values remained within the C3 diet range, we found higher *δ*^13^C during the LGM. An increase in plant *δ*^13^C has been associated with water stress (Farquhar et al. 1982; Stewart et al. 1995). Drier environmental conditions during the LGM have been documented across Eurasia by paleoenvironmental proxies (Hubberten 2004), and are further supported by bioclimatic models, which show the expansion of tundra and simultaneous reduction of boreal and temperate forests 21 ka BP (Hoogakker et al. 2016).

Post-LGM, average *δ*^13^C declined, reaching values similar to pre-LGM (Figure 3). Plants from wet environments display lower *δ*^13^C than those from dry environments (Wooller et al. 2007), and our findings of an overall decline in *δ*^13^C after the LGM may reflect an increase in moisture due to precipitation and degrading permafrost during the Late Glacial (Rabanus-Wallace et al. 2017).

We identified significant changes in *δ*^13^C over time for three of the four geographic regions (excluding W/C Europe, Figure 4A). In the E Europe/Urals and S/C Siberia, we observed a decrease in *δ*^13^C between LGM and post-LGM. In NE Siberia, we detected an increase in *δ*^13^C from pre-LGM to LGM, which remained stable during the post-LGM.

Woolly rhinoceroses ranged across the Eurasian mammoth steppe, a heterogeneous environment encompassing a mosaic of varied local environments (Szpak et al. 2010). This was reflected in our pairwise comparisons between regions, which indicated spatial differences in *δ*^13^C (Supplementary Table 4); for instance, we observed lower average *δ*^13^C in woolly rhinoceroses from NE Siberia compared with the three other regions, in particular compared to E Europe/Urals and S/C Siberia during pre-LGM and LGM (Figure 4A, Table 1). Lower *δ*^13^C has previously been associated with lower annual temperatures (Szpak et al. 2010), and this may also be the case for NE Siberia. However, this finding is unexpected, as NE Siberia was characterized by consistently arid conditions throughout the Late Pleistocene (Sher et al. 2005), and plants from dry environments have higher *δ*^13^C (Wooller et al. 2007), which should be reflected in herbivores’ *δ*^13^C compositions.

### 4.2 Temporal variation in δ^15^N among regions

Our range-wide dataset showed significant changes in *δ*^15^N over time. A significant, gradual decline in *δ*^15^N was observed across the three time bins in two of the geographic regions (W/C Europe and E Europe/Urals, Figure 4B). The decline observed from pre-LGM to LGM in S/C Siberia was not significant. A similar pattern of continuous decline in *δ*^15^N from the Late Pleistocene until after the LGM has been observed in European reindeer, attributed to the climatic cooling that culminated during the LGM (Stevens et al. 2008). The post LGM decline in *δ*^15^N, which is also reported in reindeer, has been linked to an increase in moisture due to precipitation and degrading permafrost (Drucker et al. 2003, 2011; Stevens et al. 2008). These more recent environmental changes have been attributed a major role in the Late Pleistocene and Holocene megafaunal extinctions (Rabanus-Wallace et al. 2017; Drucker et al. 2018).

Between regions, we observed some temporal differences in *δ*^15^N (Figure 4B): pre-LGM, average *δ*^15^N in NE Siberia was significantly higher than in W/C Europe and E Europe/Urals; LGM and post-LGM, the decline in average *δ*^15^N observed in other regions was not observed in NE Siberia, which continued to have high average *δ*^15^N.

Regional differences in *δ*^15^N may be driven by environmental and vegetational factors. For example, water stress and moisture influence nitrogen isotopic composition, increasing and lowering *δ*^15^N, respectively. Differences in *δ*^15^N composition have also been identified based on the ecosystem where plants grow; for instance, grasses from heath, fellfield, and boreal forest have distinct *δ*^15^N signatures (Bocherens 2003). Thus, it is not unexpected that we find differences in woolly rhinoceroses *δ*^15^N among regions, which were heterogeneous (Szpak et al. 2010). Moreover, the forms and concentrations of nitrogen that are present in the soil (i.e., mineralized or organic N) will influence the mechanisms by which plants obtain nitrogen for growth. Specifically, the interactions that plants form with symbiotic fungi (mycorrhizae) are significant, as different mycorrhizal types have varying capacities to convert organic N into mineralized N, and these capacities are strongly correlated with *δ*^15^N values in both the fungi and their plant partners (Hobbie and Högberg 2012).

The continuous decline in average *δ*^15^N observed in W/C Europe and E Europe/Urals may reflect changes in the nutrient status of the soil, such as increasing levels of recalcitrant organic nitrogen and decreasing levels of mineralized nitrogen (e.g., NH4+ and NO3-) that are readily available for plants. If the entire system becomes more dependent on fungi to process organic N and deliver it to plants, the proteolytic capabilities of those fungi impart very low *δ*^15^N values on the plants that receive the mineralized nitrogen (Hobbie and Högberg 2012).

The reduction in the number of herbivores in the landscape may have also played a role in the decline in *δ*^15^N values. Herbivores are effective recyclers of nitrogen, by depositing it back into the soil as urine and feces. An increase in the abundance of herbivores would lead to more rapid nitrogen cycling, providing more mineralized nitrogen available for plants (McNaughton 1979). Thus, there could have been an important feedback loop that was being altered in areas where megafauna was declining.

### 4.3 Temporal stability in *δ*^15^N in NE Siberia

The fossil record shows the continuous presence of woolly rhinoceroses in NE Siberia from > 50 cal kyr cal BP until their extincion ~14 cal kyr BP (Stuart and Lister 2012). The region was home to the last surviving population of woolly rhinoceros, which survived there ~1,000 years later than elsewhere.

Later persistence of open vegetation, which may have presented more suitable habitat and food resources, has been suggested to have played a role in the later survival of the species in NE Siberia (Stuart and Lister 2012). This is supported by palaeoecological proxies, which show environmental stability in NE Siberia throughout the Late Pleistocene (Clark et al. 2012; Jørgensen et al. 2012; Willerslev et al. 2014), and is further supported by our findings.

We found no significant changes in *δ*^15^N over time in NE Siberia (Figure 4B). This pattern indicates dietary stability, and that woolly rhinoceroses fed on resources with similar isotopic compositions through time. Such a scenario would require a stable environment. Temporal stability in *δ*^15^N in NE Siberia has also been observed in musk ox from Taimyr and in woolly mammoth from NE Siberia (Raghavan et al. 2014; Kuitems et al. 2019), suggesting broad-scale environmental stability, rather than a species-specific pattern.

Environmental stability may have been key to the late survival of woolly rhinoceros (and other species) in the region. In woolly mammoth, isotopic stability has been reported in both mainland NE Siberia and on Wrangel Island. Both areas were continuously inhabited by the species until their mainland and island extinctions, respectively (Arppe et al. 2019; Kuitems et al. 2019). The survival of mammoths on Wrangel Island well into the Holocene is attributed to dietary stability, and to surviving populations feeding on resources with similar isotopic composition as the Late Pleistocene mammoth population in NE Siberia (Arppe et al. 2019; Kuitems et al. 2019). We suggest a similar scenario in woolly rhinoceros, where the NE Siberian population retained a stable diet throughout the Late Pleistocene, until its extinction.

### 4.4 Isotopic niche partitioning among herbivores

In contemporaneous herbivores, interspecific differences in isotopic composition have been associated with species-specific digestion processes, dietary specialization, and niche partitioning (Drucker et al. 2003; Bocherens et al. 2015). The woolly rhinoceros is considered a grazer, primarily feeding on grass and grass-like plants (Boeskorov et al. 2011; Rivals and Lister 2016; Stefaniak et al. 2020). Morphological and genetic stomach content analysis furthermore indicates the presence of herbaceous plants (sagebrushes and forbs) (Boeskorov et al. 2011; Willerslev et al. 2014).

Overall, our results showed that woolly rhinoceros *δ*^13^C values are higher than horse and woolly mammoth, the other monogastric hindgut fermenters included in our analysis, and we suggest this may reflect ecological niche partitioning. Despite being seen as a specialized grazer, woolly rhinoceroses exhibited variable isotopic values that overlap with other large herbivores through time and space. This suggests a higher ecological flexibility than expected, and is supported by other studies on woolly rhinoceros diet (Kosintsev et al. 2019; Stefaniak et al. 2020).

We found the isotopic composition of woolly rhinoceros overlapped with musk ox and saiga, albeit during different time bins (Figure 5, Figure 6). Woolly rhinoceros pre-LGM average *δ*^13^C and *δ*^15^N differ from the other analyzed herbivores, with intermediate average *δ*^13^C values. During the LGM, woolly rhinoceros average *δ*^13^C and *δ*^15^N were similar to saiga, and post-LGM values were similar to musk ox; all three species are grazers. Saigas feed predominantly on grasses and chenopods (Hopkins et al. 2013). Musk oxen are classified as grazers based on the anatomy of their digestive system (Knott et al. 2004), and they feed primarily on grasses, sedges, and willows (Raghavan et al. 2014). The isotopic affinity of woolly rhinoceroses to these two species was not constant across time bins. Standard Ellipse Analysis shows that patterns of variation in isotopic range across time bins is species-specific (Figure 5). Thus, it is expected that isotopic niches changed across time periods, reflecting species-specific responses to environmental change.

Woolly rhinoceroses did not overlap with woolly mammoths, which may suggest ecological niche partitioning or even dietary competition between the species. Such dietary competition, and at times ecological exclusion, has been demonstrated in ecological studies of Asiatic elephants and Indian rhinoceroses (Pradhan et al. 2008), and of African elephants and black rhinoceroses (Landman et al. 2013).

### 4.5 mtDNA shows absence of phylogeographic structuring and LGM lineage expansions

Previous studies have used mtDNA to address the phylogenetic placement of woolly rhinoceros relative to other rhino species (Orlando et al. 2003; Willerslev et al. 2009), to explore the mitogenomics of NE Siberian woolly rhinoceroses (Lord et al. 2020), and to elucidate the demographic history of the species (Lorenzen et al. 2011; Lord et al. 2020). However, a comprehensive phylogeographic study across time and space has so far been lacking. To investigate spatial and temporal patterns of genetic variation across Eurasia, we compiled and analysed 61 radiocarbon dated DNA sequences from Late Pleistocene woolly rhinoceroses.

We identified three well-differentiated mtDNA haplogroups (A-C) (Figure 7). Despite the haplotype network analysis separating haplogroups A and B by only two nucleotide differences, our BEAST analysis showed strong posterior support differentiating them (Supplementary Figure 1). Our analysis included six control region sequences from Lord et al. (2020), which in that study, based on entire mitochondrial genomes, were assigned to woolly rhinoceros mitogenome clade 1 (n = 2) and clade 2 (n = 4). Mitogenome clade 1 corresponded to our haplogroup B, and mitogenome clade 2 corresponded to our haplogroup A, further supporting our haplogroup division based on the shorter control region sequences.

The mtDNA control region haplogroups were not geographically structured in time or space, and extended from Central Europe to NE Siberia; the only mtDNA sequence included from W/C Europe grouped in haplogroup C (Figure 7). During the pre-LGM, haplogroup A was present in NE Siberia only, with an LGM expansion into S/C Siberia; the first appearance of haplogroup A in S/C Siberia was dated to 26,030 cal yr BP. A similar pattern was observed in haplogroup B; pre-LGM it was limited to S/C and NE Siberia, and expanded into E Europe/Urals during the LGM. The expansion of haplogroups A and B into new regions occurred at a time where demographic analysis of subsets of the data indicate an increase in population size (Lorenzen et al. 2011; Lord et al. 2020).

The presence of spatially overlapping yet divergent haplogroups likely reflects the allopatric divergence of lineages, with subsequent expansion leading to the distribution of woolly rhinoceros populations across northern Eurasia. Siberia harbored the highest levels of genetic diversity, and was the only area where all three haplogroups were found. However, DNA analysis of additional specimens from the western part of the species range are needed to confirm this pattern.

For other megaherbivore species, such as bison, high genetic diversity has also been identified in NE Siberia (Kirillova et al. 2015a; Massilani et al. 2016). Furthermore, genetic data from wolves (*Canis lupus*) suggest Late Pleistocene populations likely originated and expanded from NE Siberia into Eurasia and North America (Loog et al. 2020). Climatic stability has been suggested as a possible explanation for this pattern of maintained genetic diversity in NE Siberia (Loog et al. 2019); a scenario supported by our study.

Demographic analysis of 55 of the 61 mtDNA sequences analyzed here showed changes in genetic connectivity through time, with increasing fragmentation prior to extinction leading to population isolation (Lorenzen et al. 2011). The phylogeographic pattern revealed in our analysis supports this hypothesis of a decline in connectivity over time; we observed some changes in the geographic distribution of haplogroups over time, although we did not detect a loss of mtDNA haplogroups from the pre-LGM to post-LGM. Two of the youngest samples included in our study (dated to 15,620 and 14,390 cal yr BP) were from NE Siberia and represented haplogroups A and B, respectively (Figure 7). The youngest haplogroup C sample was from E Europe/Urals and is dated to 16,620 cal yr BP.

Previous analysis of 14 mitochondrial genomes from NE Siberia also support lineage continuity from the pre-LGM to the post-LGM in this region (Lord et al. 2020). However, further data are needed to confirm this pattern, as inferences in (Lord et al. 2020) were based on a small number of individuals, with an overrepresentation of sequences from NE Siberia.

## 5. Implications for woolly rhinoceros extinction

We investigated a comprehensive *δ*^13^C and *δ*^15^N dataset of woolly rhinoceros, and included the first 71 stable isotope records spanning the 15,000 years preceding species extinction. Our study underscores the applicability of combining range-wide stable isotope studies with ancient DNA analysis, to obtain a broader understanding of the evolutionary ecology of past populations. We uncovered ecological flexibility and geographical variation in woolly rhinoceros, and suggest spatial and temporal variation in isotopic compositions were driven by environmental and vegetational factors. Our analysis showed Late Pleistocene stability in *δ*^15^N in NE Siberia, which we suggest reflects long-term environmental stability that may have favoured the later survival of woolly rhinoceros in the region.

Our comparative analysis with contemporaneous herbivores suggested possible niche partitioning of woolly rhinoceros. We detected temporal variation in woolly rhinoceros’ isotopic profile compared with other herbivores, which further supports the ecological flexibility of the species.

Ancient DNA analyses showed a lack of geographical structure at the mitochondrial level, with divergent lineages overlapping in time and space, as has been shown with a more spatially and temporally limited dataset from NE Siberia (Lord et al. 2020). We did not detect lineage loss prior to species extinction.

Stable isotope analyses have been used to investigate causes of extinction in other herbivores. For woolly mammoth, isotopic evidence suggested either a population decline (due to human encroachment) allowing other species (horse) to occupy their niche (Drucker et al. 2015), or a niche change forced by climate-induced environmental change (Drucker et al. 2018). Our data did not support either of these scenarios for the woolly rhinoceros.

Radiocarbon chronologies and genetic data have correlated the extinction of woolly rhinoceroses with climatic and environmental factors (Lorenzen et al. 2011; Lister and Stuart 2012; Lord et al. 2020). Genetic analysis indicated an increase in population size ~30 cal kyr BP followed by demographic stability until ~18.5 cal kyr BP, where a decline in woolly rhinoceros population size started. Our results indicated that environmental stability enabled the late survival of woolly rhinoceros in NE Siberia, and thus, environmental changes around the time of extinction may have had detrimental effects on the remaining populations. Pollen records for the region showed a spread of shrub tundra communities and increased precipitation between 13.5 - 12.7 cal kyr BP, compared to the open steppe-tundra community that was previously found in the region (until ~15 cal kyr BP) (Müller et al. 2008). We suggest the interplay between environmental instability and fragmentation/isolation of populations, as presented in Lorenzen et al. (2011), played a major role in the extinction of the woolly rhinoceros.

## Authors Contributions

ARI: Data Curation, Formal analysis, Visualization, Writing - Original Draft. AML: Resources, Funding acquisition, Writing - Review & Editing. AJS: Resources, Funding acquisition. HB: Conceptualization, Formal analysis, Writing - Original Draft. PS: Writing - Original Draft, Conceptualization. EW: Resources, Funding acquisition, Writing - Review & Editing. EDL: Conceptualization, Funding acquisition, Writing - Original Draft.

