## Supplemental Figures and Supplemental Tables 4-6 for "Late Pleistocene palaeoecology and phylogeography of woolly rhinoceroses"

**Supplementary Figures and Tables:**

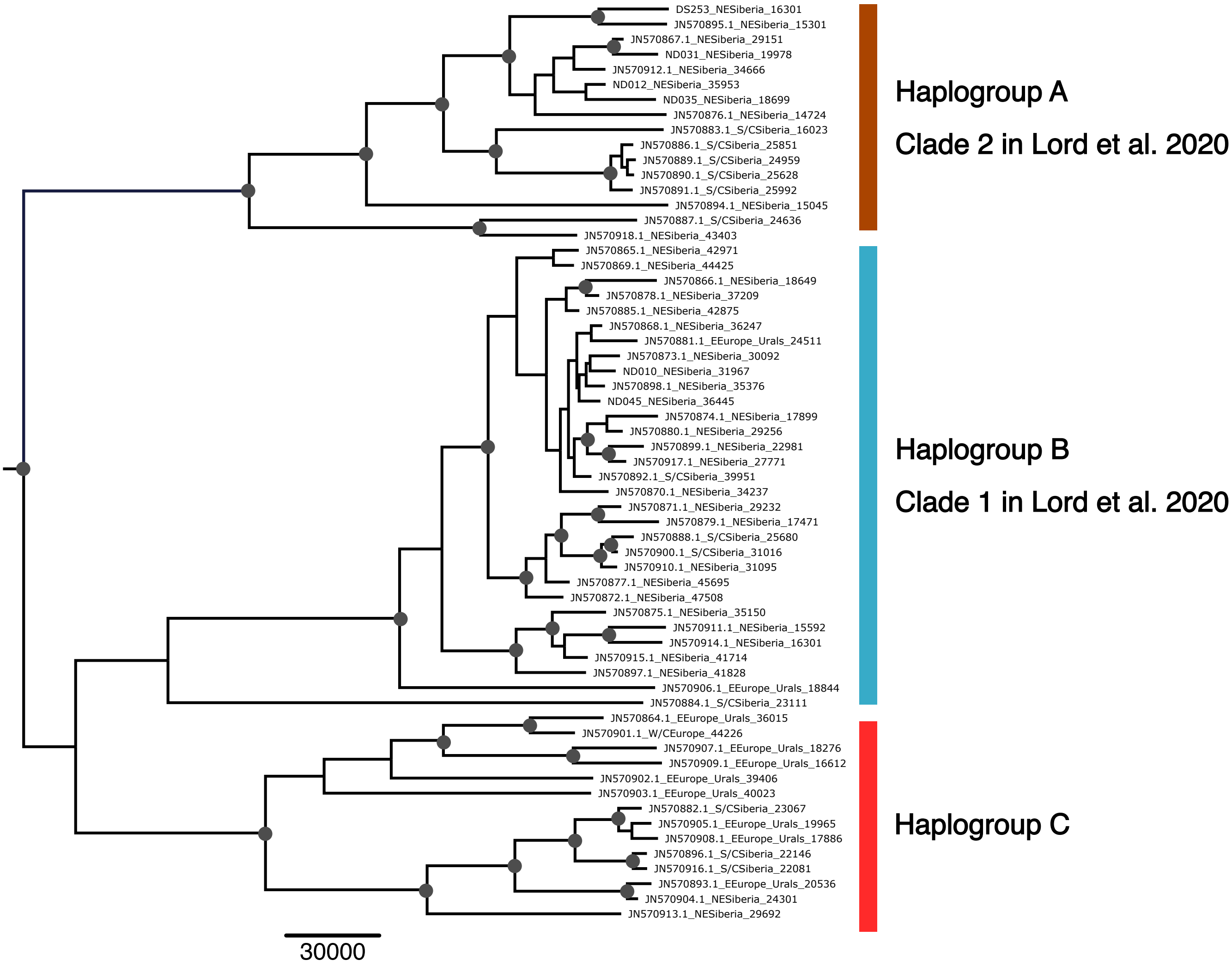


**Supplementary Figure 1.** Tip-calibrated phylogeny of 61 woolly rhinoceros mtDNA control region sequences using BEAST. Red nodes indicate posterior probabilities > 0.65. Tip label postfixes indicate sample region and age. Coloured bars represent the haplogroups identified in the network analysis (Figure 7).

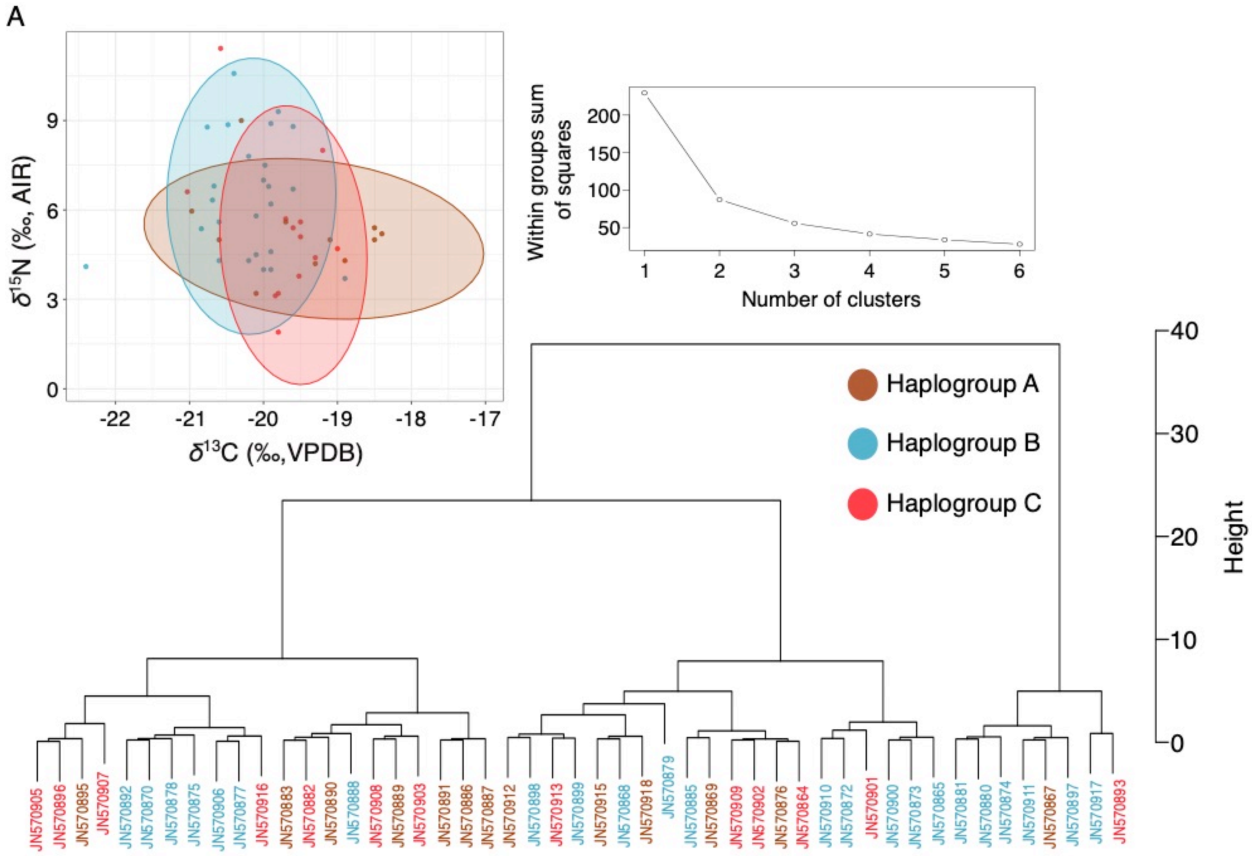


**Supplementary Figure 2.** Bivariate plot and cluster analysis of $\delta^{13}$C and $\delta^{15}$N in the 49 woolly rhinoceros included in this study for which we also have mitochondrial data. (A) Bivariate plot of $\delta^{13}$C and $\delta^{15}$N. Ellipses represent 0.95 confidence levels. (B) Cluster analysis using $\delta^{13}$C and $\delta^{15}$N. In both plots, colours represent the mitochondrial haplogroup for each of the 49 samples.

**Supplementary Table 1.** Summary table of the 286 published and unpublished woolly rhinoceros samples used in the $\delta^{13}$C and $\delta^{15}$N analysis.

**Supplementary Table 2.** Summary table of the five coeval mammoth steppe herbivores used for comparison in this study

**Supplementary Table 3.** List of the 61 published mitochondrial sequences analyzed in this study.

**Supplementary Table 4.** Statistical comparison to identify temporal differences in the isotopic composition of woolly rhinoceros between the four regions explored in this study, using the non-parametric Kruskal Wallis and the post hoc Kruskal Conover tests. Asterisk indicates statistically significant differences

| **Time bin** | **Region** | ***δ*13C** | | | | ***δ*15N** | | | |
|---|---|---|---|---|---|---|---|---|---|
| | | W/C Europe | E Europe /Urals | S/C Siberia | NE Siberia | W/C Europe | E Europe /Urals | S/C Siberia | NE Siberia |
| pre-LGM (>24,600 yr BP / 28,660 cal yr BP) | W/C Europe | - | - | - | - | 0.99 | - | - | - |
| | E Europe/Urals | < 0.001* | - | - | - | 0.99 | - | - | - |
| | S/C Siberia | 0.010 | 0.176 | - | - | 0.13 | 0.19 | - | - |
| | NE Siberia | < 0.001* | < 0.001* | < 0.001* | - | < 0.001* | 0.0034* | 0.59 | - |
| LGM (24,600-17,000 yr BP / 28,660-20,520 cal yr BP) | W/C Europe | - | - | - | - | | - | - | - |
| | E Europe/Urals | 0.47 | - | - | - | 1.00 | - | - | - |
| | S/C Siberia | 0.47 | 0.94 | - | - | 1.00 | 1.00 | - | - |
| | NE Siberia | 0.76 | 0.026* | 0.007* | - | 0.75 | 1.00 | 1.00 | - |
| post-LGM (<17,000 yr BP / < 20,520 cal yr BP to extinction) | W/C Europe | - | - | - | - | - | - | - | - |
| | E Europe/Urals | 1.00 | - | - | - | 1.00 | - | - | - |
| | S/C Siberia | 1.00 | 1.00 | - | - | 0.22 | 0.19 | - | - |
| | NE Siberia | 1.00 | 0.14 | 1.00 | - | 0.19 | 0.035* | 1.0 | - |

**Supplementary Table 5.** Summary table of the statistical comparisons carried out to identify dietary differences in isotopic composition between woolly rhinoceros and the five coeval megaherbivores surveyed. Data was divided by time bin, and comparisons were performed using the non-parametric Kruskal Wallis and the post hoc Kruskal Conover tests.

| | | ***δ*13C** | ***δ*15N** |
|---|---|---|---|
| **Time Bin** | **Species** | Woolly rhinoceros | Woolly rhinoceros |
| pre-LGM | Horse | < 0.001 | 0.98 |
| | Musk ox | < 0.001 | 0.98 |
| | Reindeer | < 0.001 | < 0.001 |
| | Saiga | < 0.001 | < 0.001 |
| | Woolly mammoth | < 0.001 | < 0.001 |
| LGM | Horse | < 0.001 | < 0.001 |
| | Musk ox | < 0.001 | 0.002 |
| | Reindeer | 1.00 | < 0.001 |
| | Saiga | 1.00 | 0.19 |
| | Woolly mammoth | < 0.001 | < 0.001 |
| post-LGM | Horse | < 0.001 | < 0.001 |
| | Musk ox | 0.90 | 0.61 |
| | Reindeer | < 0.001 | < 0.001 |
| | Saiga | < 0.001 | 0.73 |
| | Woolly mammoth | < 0.001 | < 0.001 |

**Supplementary Table 6.** Summary table of the Standard Ellipse Area values estimated using SIBER for each species and time bin.

| **Species / Time bin** | Horse | Musk ox | Reindeer | Saiga | Woolly mammoth | Woolly rhinoceros |
|---|---|---|---|---|---|---|
| pre-LGM | 2.58 | 2.85 | 1.27 | 5.16 | 2.30 | 4.25 |
| LGM | 0.48 | 1.36 | 1.20 | 5.32 | 2.91 | 3.65 |
| post-LGM | 3.00 | 3.00 | 1.17 | 1.11 | 4.13 | 4.98 |
